# Applying multi-state modeling using AlphaFold2 for kinases and its application for ensemble screening

**DOI:** 10.1101/2024.04.04.588044

**Authors:** Jinung Song, Junsu Ha, Juyong Lee, Junsu Ko, Woong-Hee Shin

## Abstract

Structure-based virtual screening (SBVS) is a pivotal computational approach in drug discovery, enabling the identification of potential drug candidates within vast chemical libraries by predicting their interactions with target proteins. The SBVS relies on the receptor protein structures, making it sensitive to structural variations. Kinase, one of the major drug targets, is known as one of the typical examples of an active site conformation change caused by the type of binding inhibitors. Examination of human kinase structures shows that the majority of conformations have the DFGin state. Thus, SBVS using the structures might cause a favor of type of ligand type I inhibitors, bind to the DFGin state, rather than finding the diverse scaffolds. Recent advances in protein structure prediction, such as AlphaFold2 (AF2), offer promising solutions but may still be possibly influenced by the structural bias in existing templates. To address these challenges, we introduce a multi-state modeling (MSM) protocol for kinase structures. We apply MSM to AF2 by providing state-specific templates, allowing us to overcome structural biases and thus apply them to kinase SBVS. We benchmarked our MSM models in three categories: quality of predicted models, reproducibility of ligand binding poses, and identification of hit compounds by ensemble SBVS. The results demonstrate that MSM-generated models exhibit comparable or improved structural accuracy compared to standard AF2 models. We also show that MSM models enhance the accuracy of cognate docking, effectively capturing the interactions between kinases and their ligands.

In virtual screening experiments using DUD-E compound libraries, our MSM approach consistently outperforms standard AF2 modeling. Notably, MSM-based ensemble screening excels in identifying diverse hit compounds for kinases with structurally diverse active sites, surpassing standard AF2 models. We highlight the potential of MSM in broadening the scope of kinase inhibitor discovery by facilitating the identification of chemically diverse inhibitors.

**Author Summary:** One of the main problems with structure-based virtual screening is structural flexibility. Ensemble screening is one of the conventional approaches to solving the issue. Gathering experimental structures or molecular simulations could be used to compile the receptor structures. Recent developments in algorithms for predicting protein structures, like AlphaFold2, suggest that different receptor conformations could be produced. However, the prediction approaches produce biased structures because of the bias in the structure database. In order to solve the problem, we developed a protocol called multi-state modeling for kinases. Rather than supplying multiple sequence alignments as an input, we gave the AlphaFold2 a specific template structure and the sequence alignment between the template and query.

Our findings imply that our technique can yield a particular structural state of interest with an enhanced or comparable structural quality to AlphaFold2 and predict highly accurate protein-ligand complex structures. Lastly, compared to the typical AlphaFold2 models, ensemble screening using the multi-state modeling approach improves the structure-based virtual screening performance, particularly for diverse active molecular scaffolds.

## Introduction

Structure-based virtual screening (SBVS) is one of the most widely used computational drug discovery approaches to identify novel active compounds against the target from a virtual molecule library. By predicting the interaction between the ligand and the target protein, the technique ranks the ligands based on their scores that mimic the binding affinity between the molecules. The SBVS is a cost- and time-efficient way to explore the vast chemical space, by significantly narrowing down the number of drug candidates that need to be synthesized and tested in experiments. Calculating the interaction between the receptor and ligand can be done in various ways: molecular docking, molecular dynamics, fingerprint, pharmacophore matching, and so forth. As its name implies, the method requires the target protein structure, and the performance depends on the protein conformation. For targets whose experimental structures are not available, researchers should predict the three-dimensional structures using methods such as homology modeling.

One of the major obstacles in the SBVS method, especially molecular docking, is caused by its static picture of a receptor structure. Proteins are flexible molecules, so they can change their shape depending on the binding partners. The structural change for receptor protein might lead to a failure of molecular docking [1–3]. One of the techniques for treating receptor flexibility is ensemble screening. The method uses pre-generated diverse receptor structures gathered from experimental structure databases or simulation trajectories such as molecular dynamics or normal mode analysis. Ligands in the screening library are docked to individual structures and ranked by their representative scores. There are various methods to get the representative score: arithmetic mean, harmonic mean, etc. Since the ensemble method uses multiple target structures, it is important to reflect the structural diversity when selecting the receptor ensemble. However, the crystal structures of the target would be biased to thermodynamically stable states or the major type of inhibitor-bound form. This might make the screening for obtaining diverse scaffolds or even lead to a failure in the SBVS even structural ensembles are used.

Kinases are one of the typical examples having structural diversity. They play a key role in biological processes in phosphorylation transferring a phosphate group from ATP to proteins. Since the process modulates key cellular operations like cell cycle regulation, metabolism, and apoptosis, kinases have become attractive targets in drug discovery. According to Santos et al. [4], kinases belong to one of the four main target families, which are G protein-coupled receptors (GPCRs), ion channels, nuclear receptors, and kinases. In the ChEMBL database [5], the kinases are targeted by 14.2% of compounds.

The kinase domain, a structural domain with catalytic function, has a highly conserved structure. It is composed of an N-lobe and a C-lobe, linked by a hinge region [6]. The N-lobe is structured with five β-strands and a single alpha helix, known as the C-helix, while the C-lobe is comprised of multiple alpha helices. The catalytic activity of kinase domains is regulated by two loops, namely the activation loop and the catalytic loop. The His-Arg-Asp (HRD) motif of the catalytic loop directly interacts with the hydroxyl group of the target protein residue (serine, threonine, or tyrosine) set for phosphorylation. The Asp-Phe-Gly (DFG) motif found in the N-terminal of the activation loop plays a crucial role in anchoring ATP to the active site. The conformational states of the active site of kinase domains are classified as DFGin, DFGinter, or DFGout, based on the orientation of the aspartic acid of the DFG motif relative to the site. Fig 1 illustrates two distinct structural states, DFGin (Fig 1A) and DFGout (Fig 1B), of BRAF. The DFGin state locates Phe of the DFG motif into the ATP binding pocket, allowing kinase to hold ATP, thus it is called the active state. On the other hand, DFGout conformation directs the Phe out of the ATP binding pocket, so it is classified as an inactive state.

**Fig 1.**
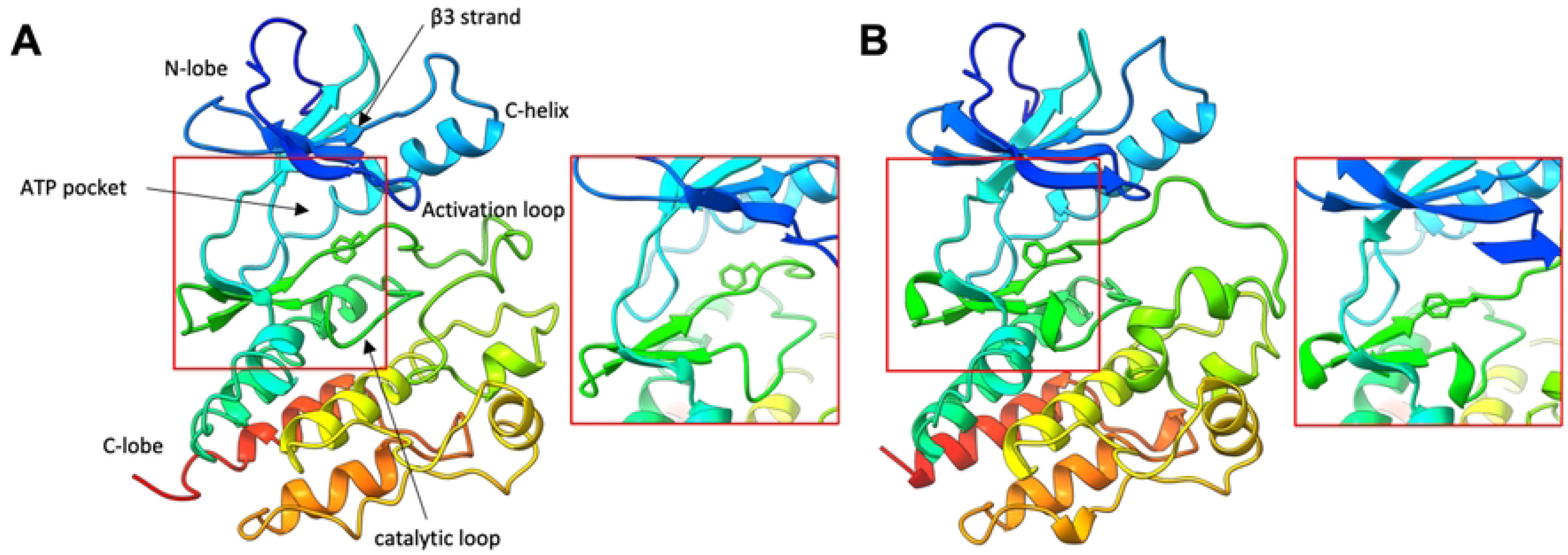
The structure of kinase domain and key structural elements. The active state (DFGin-BLAminus) of BRAF (PDB ID: 6UAN, **A**) and the inactive state (DFGout-BBAminus) of BRAF (PDB ID: 8C7X, **B**). The active site is focused in the red box on the right of each panel. The activation loop and DFG motif changes conformation depending on the state.

The kinase inhibitors can be categorized depending on the binding site conformation of kinases and the place the compounds bind. Type I inhibitors are the ones that bind to the ATP-binding site and thus compete with ATP. Most type I inhibitors bind to the DFGin state. We observed that the majority of experimentally determined human kinase structures form DFGin state (87%, as of May 2023). On the other hand, type II inhibitors often associate with the kinase to the DFGout state. The type II compounds tend to occupy ATP-binding site partially and a hydrophobic pocket close to the ATP-binding site, which is opened when the activation loop forms the DFGout conformation. In general, type II inhibitors have more selectivity than type I inhibitors. According to Hari et al. [7], the type II inhibitors’ selectivity is influenced by the inherent variations in kinase’s capacity to adopt the DFGout conformation. Lastly, type III inhibitors, sometimes referred to as allosteric inhibitors, bind to a kinase site that is not directly connected to the ATP-binding site. Therefore, it is crucial to take into account as many different structural states of kinases as possible to find diverse hit molecules for SBVS.

Recent advances in protein structure prediction using deep learning techniques, such as AlphaFold2 (AF2) [8] and RoseTTAFold [9], allow for accurate modeling of protein structures. However, these methods rely on pre-trained models from the PDB database, so the generated models might be affected by the conformational state distribution in the PDB if the protein can form diverse structures. Therefore, the methods could produce kinase structures similar to the DFGin state, and thus SBVS targeting kinases using the predicted models could potentially yield outcomes favoring type I inhibitors even using the modeled structures.

To solve the issue, Heo and Feig [10] suggest a method called multi-state modeling (MSM) to predict GPCRs with high accuracy of the desired structural state. The authors found that predicted models by AF2 tend to have either active or inactive states depending on GPCR classes due to the small number of experimentally determined structures of GPCRs. Instead of providing multiple sequence alignment (MSA) as an input of AF2, the authors align a query sequence to a template sequence with the structural state of interest. The MSM technique showed an improved performance of modeling for GPCRs. Cognate docking using MSM models also showed enhanced accuracy with smaller root-mean-square-distance (RMSD) from the crystal structure.

Inspired by the work, we established an MSM protocol for modeling the kinase structures to overcome the structural bias of kinases for SBVS by giving state-specific templates to AF2. All human kinase experimental structures were classified by the active site conformation using KinCoRe [11] to construct a state-specific template database. Our protocol was able to predict kinase conformations with the desired structural state with a high accuracy. Then we benchmarked cognate docking to observe the MSM models produced accurate binding modes with lower RMSD than standard AF2 models. Finally, we performed ensemble SBVS with generated models by MSM. The ensemble method showed higher performance compared to screening with modeled structures using AF2, especially when the active molecules are diverse.

## Results and Discussion

### Conformation Distributions of Experimental Kinase Structures and AlphaFold2 Predicted Models

Proteins are flexible molecules, so they may change their structures when bind to their partners. This conformational change could act as an obstacle to SBVS. The active site of kinases also forms diverse conformations depending on the type of the binding compound. For example, the DFGin state binds to the type I inhibitor, while the DFGout state binds to the type II inhibitor. Dunbrack and his colleagues introduced a rule, called KinCoRe [11], to classify the kinase structure. The rule classifies structural states of kinase structures into 12 categories based on the spatial state of the activation loop and the dihedral angle of DFG motif. Details of the criteria can be found in the Materials and Methods section and Modi et al [11].

Fig 2 shows a distribution of kinase conformational states annotated by KinCoRe scheme [11] in the PDB database (blue bar). Based on the KinCoRe notation, more than half (53.6%) of experimentally determined kinase structures have DFGin-BLAminus conformation. Details of the structure distribution are shown in Supplementary Material (S1 Table). Other than the major state, the other conformational states occupy less than 10% of PDB structures.

**Fig 2.**
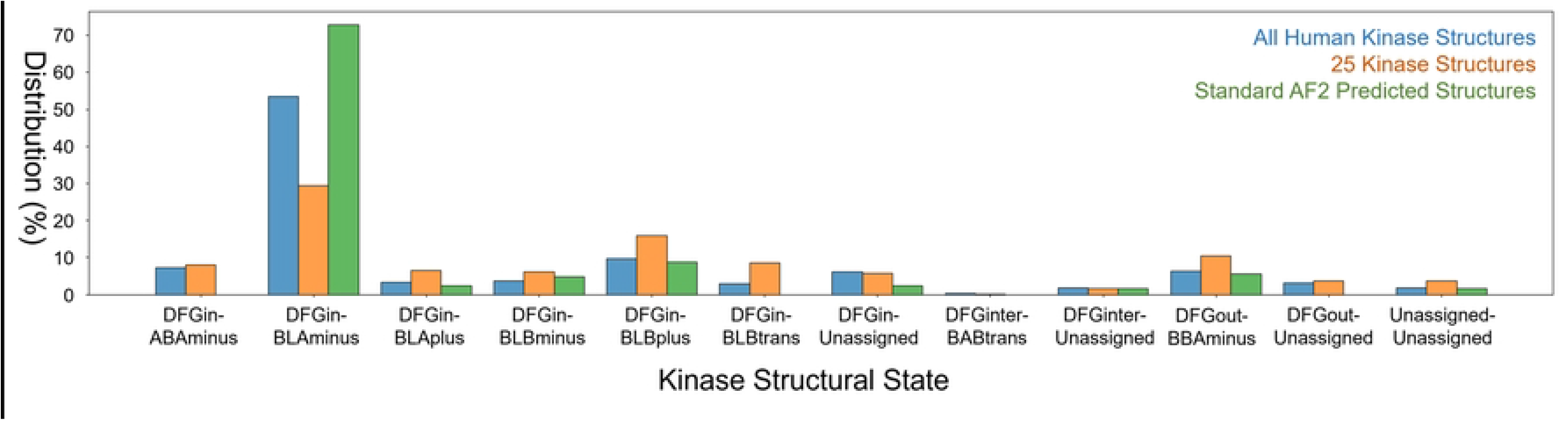
Distribution of crystal structures and standard AF2 structures for each kinase conformation. The X-axis is conformational states annotated by KinCoRe and the y-axis is the percentage of each state. All human crystal kinase structures deposited in RCSB-PDB, DUD-E target protein crystal structures, and predicted AF2 models with default parameters (standard AF2) are colored as blue, orange, and green, respectively.

The preference for the DFGin state in experimentally determined structures might be influenced by the thermodynamic stabilities of the conformational states. Meng et al [12], studied the transition between the structural states of c-Abl and c-Src kinases using umbrella sampling and the potential of mean force. With a difference of 1.4 kcal/mol, the DFGin state of c-Abl has a lower Gibbs free energy than the DFGout state. The calculation shows the active conformation (DFGin) is the dominant population while the DFGout state occupied only 9% in their simulation. Similarly, for c-Src, the DFGin state is more favored than the DFGout state. The free energy difference between the states is calculated as 5.4 kcal/mol. The authors also studied a thermodynamic barrier of the transition between the states [13]. It is estimated as the order of 2-4 kcal/mol making the transition from highly populated the DFGin state to the DFGout state hard. Levy and his colleagues [14] collected 2,896 kinase structures and multiple sequence alignment and applied Potts model to predict structural propensities from sequences. With the statistical potential, most of the kinases are predicted to have preferences for the DFGin state. The penalty for forming the DFGout state reaches 2-3 kcal/mol for the extreme case.

Not only for the thermodynamic preference, but the type of kinase inhibitor distribution might also affect the skewness of kinase structures. We counted the number of compounds in kinase inhibitor types from PKIDB (assessed August 2023) [15]. PKIDB is a curated database of kinase inhibitors in clinical trials. Out of 369 compounds in the database, only 84 molecules have their inhibition type annotation, because annotating the ligand type needs the complex structure. Type I inhibitor (DFGin bound) is the dominant form, occupying 66% (55 compounds) of the annotated inhibitors. In contrast, 17 molecules are labeled as type II (DFGout bound). Thus, DFGin conformation (active state) might have more chance to be crystallized than the DFGout state. The biased conformational states of kinases would make the discovery of chemically diverse kinase inhibitors harder.

We also examined the conformational state distribution of 25 DUD-E targets [16], that are benchmarked throughout this study, in the PDB database (orange bar). DUD-E is a widely used benchmark dataset for evaluating virtual screening methods, composed of pharmaceutically important targets such as GPCRs, kinases, and nuclear receptors. Among the set, we extracted 25 kinase targets for benchmarking. The experimental structures are less biased than all PDB structures, about 30% of DUD-E proteins have DFGin-BLAminus conformation. The other states tend to have a higher proportion than the PDB structure distribution of the states. The difference in the state distribution between all kinases and DUD-E targets potentially means that the highly biased nature of kinase structures might not be suitable for finding diverse types of kinase inhibitor discovery.

From an MSA of a given sequence, AF2 [8] extracts coevolution information by the MSA Transformer and predicts the three-dimensional structure based on the information and deep-learning models trained on existing protein structures. We modeled 25 kinase catalytic domains provided in the DUD-E kinase subset with the default parameters of AF2 resulting in 125 structures (five models per target), then assigned the conformational states of the models using KinCoRe. Throughout this paper, AF2 with default parameters is called standard AF2. The average plDDT and MolProbity [17] score of the predicted structures is 89.38 and 1.04, respectively. This implies that they are properly modeled. Out of 125 predicted models, 91 structures (72%) are annotated to have DFGin-BLAminus conformation (green bar). The distribution of predicted models is more skewed than all human kinase structures and the DUD-E set proteins. However, the other states have a lower proportion than the experimental structures.

The standard AF2 did not produce DFGin-ABAminus, DFGin-BLBtrans, DFGinter-BABtrans, and DFGout-Unassigned conformations, which occupy a small portion of the human kinase PDB structures (7.3%, 3.0%, 0.3%, and 3.2%, respectively, S1 Table). Since the experimentally determined kinase structures are biased to the DFGin-BLAminus, predicted kinase structures by AF2 might have a chance to be biased to the conformation, which can be observed in previous research [10]. In addition to the biased trained models, the template selection process in AF2 modeling does not take the structural state of the kinase into account.

### Predicting State-specific Kinase Structures using Multi-state Modeling Protocol

AF2 with MSM protocol provides conformational state-specific structures as templates for AF2 models. All human kinase structures from the KLIFS database [18] and further classified by their structural state following KinCoRe rules [11]. For each state, the five highest sequence similar structures were selected as templates. Then AF2 was executed with the template information to predict five models for each template. The models with undesired conformational state were removed, and then the model with the highest plDDT was selected for our benchmark. The overall workflow is illustrated in Fig 3. Details of the kinase MSM protocol are elaborated in the Materials and Methods section.

**Fig 3.**
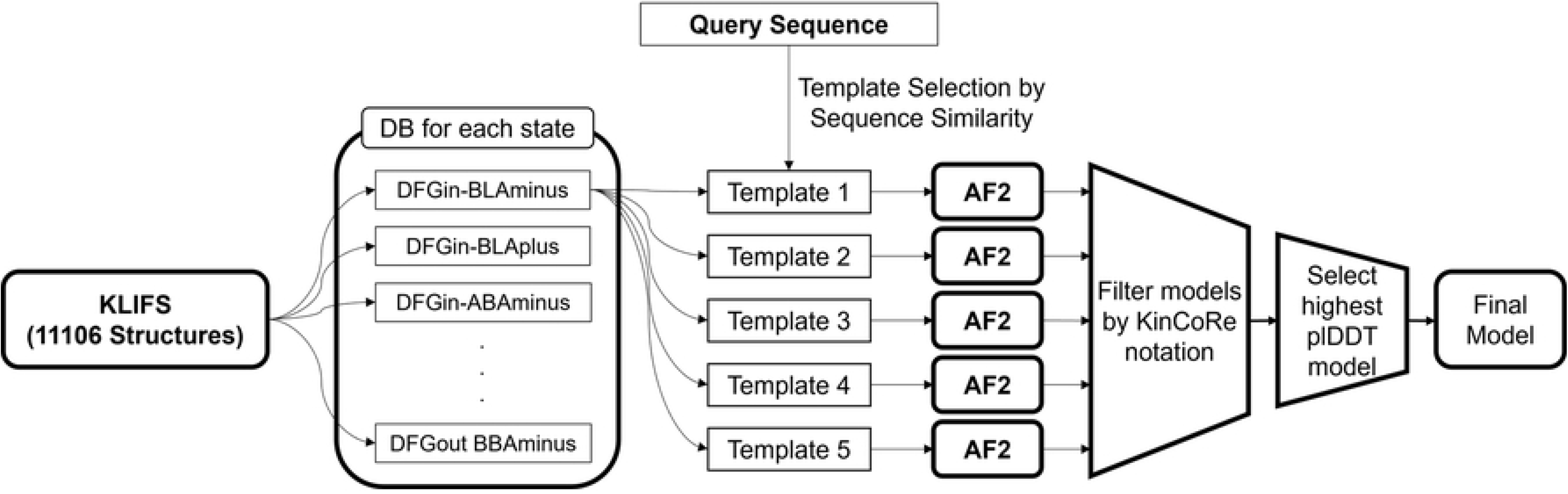
Workflow of multi-state modeling of kinases.

We prepared two template sets, a trivial template (TT) set containing 100% sequence identical structural template and a nontrivial template (NT) set not having identical protein. With our MSM protocol, AF2 was able to model structures in a specific state, producing 8.5 and 8.2 models with TT set and NT set on average, respectively. The number of predicted models for each target ranges from three (WEE1) to 11 (ABL1 and LCK, S2 Table). The average plDDT scores of structures are 89.63 and 88.08 for models from the TT set and NT set, respectively, showing comparable values to the standard AF2 predictions (89.38). As expected, TT set models have higher accuracy on average than NT set, but their difference is marginal. The distribution of the MolProbity score is given in S1 Fig. We also examined the quality of models for each structural state. The average plDDT values range from 86.62 (DFGin-BLAplus) to 94.48 (DFGin-BLAminus), meaning that the quality of the models does not depend much on the structural state of kinases. The highest plDDT score of DFGin-BLAminus might be caused by the highest frequency of the state in the PDB database (S1 Table). The Pearson’s correlation coefficient between the average plDDT and percentage in the PDB structure of each structural state is 0.739. Even though the MSM protocol provides a structural template for AF2, the model quality might be influenced by the pre-trained models of AF2, so the average plDDT of each model follows the distribution of the crystal structure.

The TT set model and the NT set model have average MolProbity scores of 1.21 and 1.24, respectively. These figures represent a modest decline from the baseline AF2 models (1.04). We also examined each structural state’s average MolProbity score. 1.06 (DFGin-ABAminus) is the lowest value, and 1.33 (DFGinter-Unassigned) is the highest. For DFGin-BLAminus, the highest populated states for both PDB and standard AF2, MSM models show the average MolProbity score as 1.08. The average MolProbity score and the distribution in the PDB structure have a −0.59 Pearson’s correlation coefficient, indicating a weak correlation, and the high score of MolProbity might be caused by the states with the small PDB populated states.

To investigate the accuracy of the models, we measured the TM-Score [19] of predicted models to crystal structures given in the DUD-E set, called ‘reference structure’ throughout this paper. TM-Score assesses a structural similarity between two given proteins, ranging from zero (not similar) to one (identical). For MSM models, we used the predicted structures in the same structural state as the reference. The average TM-Scores are 0.92 and 0.90 for the models predicted using TT and NT sets, respectively. Compared with standard AF2 models (average TM-Score: 0.87), the MSM technique provided more similar models than the standard AF2 protocol, which is expected since the MSM provides structural templates. Models with TT sets generally have more accurate structures than those with NT sets as also expected.

Consequently, by utilizing structures that represent a variety of states for the target kinase, the MSM is able to generate diverse structures as desired with high accuracy. Thus, it could provide a proper structure set for kinase ensemble SBVS.

### Cognate Docking Accuracy of a Compound to the Multi-state Modelled Structures

To examine whether modeled structures are suitable for molecular docking and thus structure-based virtual screening or not, we first conducted a cognate docking experiment on the predicted structures, both standard AF2 and MSM models. Ligands from complex crystal structures of the DUD-E kinase subset were used for this benchmark. For the standard AF2 model, the highest plDDT model for each protein, which is also used for performing virtual screening in the next section, was selected for evaluation. For evaluating the MSM protocol, we used the modeled structure with the same KinCoRe annotation as the reference PDB structure provided by DUD-E. Out of 25 kinases in DUD-E, IGF1R was removed, since the crystal structure was not assigned any of the structural states by KinCoRe, Therefore, the number of target proteins becomes 24. AutoDock-GPU [20] was employed to predict the 50 binding poses. The RMSDs of all predicted poses to the crystal binding pose were calculated. To analyze, we took three values: the RMSD of the best AutoDock score pose, that of the closest pose to the crystal binding mode, and the average RMSD of all 50 poses.

Table 1 summarizes the docking accuracy evaluation results. Taking the best AutoDock score models, the standard AF2, MSM structures modeled with TT, and NT have average RMSD of 2.74 Å, 2.15 Å, and 3.49 Å, respectively. In addition, the success cases with an RMSD cutoff of 2.0 Å, a standard criterion for judging docking success [21–23], are 11 (standard AF2), 16 (with TT), and 9 (with NT) out of 24 receptors. Individual RMSD values are given in S3 Table.

**Table 1.** Docking accuracy benchmark result. The values are the average values of 24 proteins and the numbers in the parentheses are the number of success cases with RMSD < 2 Å.

| Structure | Standard AF2 | MSM with TT | MSM with NT |
| --- | --- | --- | --- |
| **Best AutoDock Score** | 2.74 (11) | 2.15 (16) | 3.49 (9) |
| **Lowest RMSD** | 1.50 (20) | 1.40 (19) | 2.14 (14) |
| **Average RMSD** | 2.93 | 2.49 | 3.34 |

Comparing the MSM models and AF2 models, the multi-state models with TT sets have the most accurate docking poses. As observed in the previous section, template information influenced the quality of the docking poses, i.e., the predicted docking poses of the models with TT sets have smaller RMSD and more successful cases than those with NT sets. One successful example is AKT2 (Fig 4). Although the TM-scores of MSM with TT set (0.98) and standard AF2 (0.96) are similar, the best AutoDock score models docked to the MSM using TT sets has more accurate binding pose than standard AF2, RMSD of 0.86 Å (magenta, Fig 4B) and 3.24 Å (cyan, Fig 4C), respectively.

**Fig 4.**
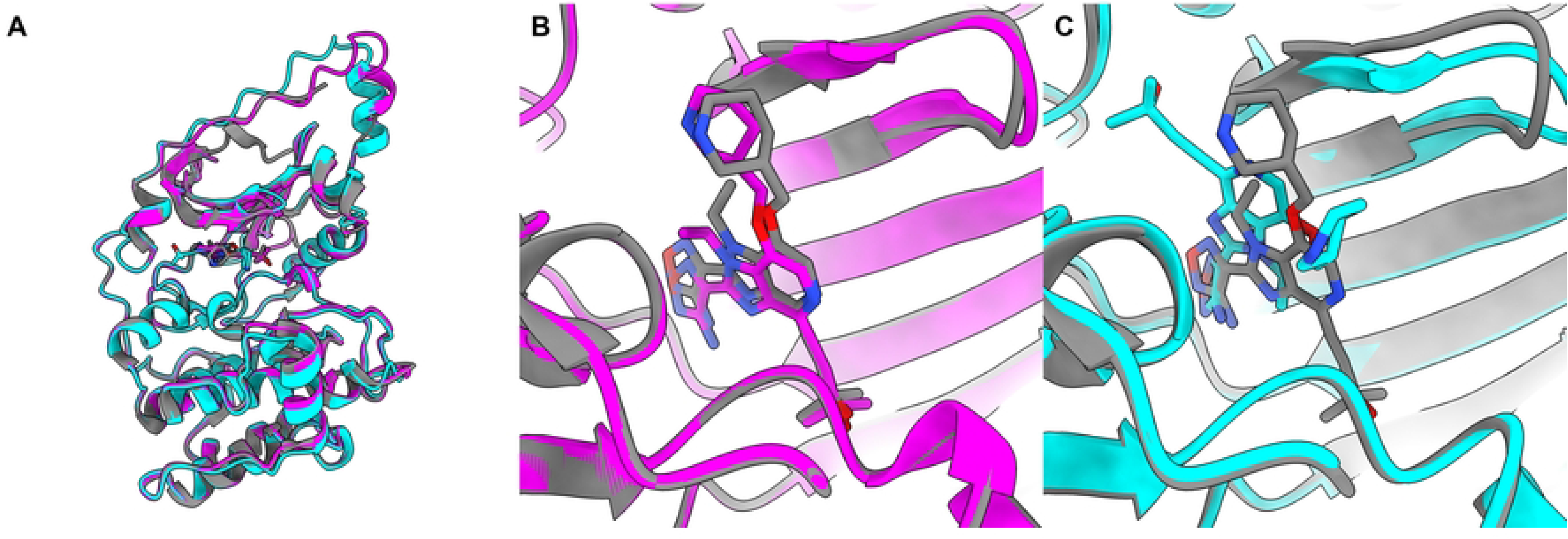
Predicted cognate docking structures of AKT2. **A.** Superimposed structures of crystal structure (PDB ID: 3D0E, grey), MSM model (DFGin-BLAminus, magenta; TM-Score: 0.98), and standard AF2 (cyan; TM-Score: 0.96). **B.** Predicted docking poses of the cognate ligand on the AF2 with MSM. The RMSD of the predicted pose is 0.86 Å. **C.** Predicted docking poses of the cognate ligand on the standard AF2. The RMSD of the predicted pose is 3.24 Å.

Although the docking is not successful (RMSD > 2 Å), MSM structure was able to retrieve the interaction between protein and ligand found in the crystal structure for some cases. Out of eight cases, the interacting residues in the reference structures were successfully captured more than 50% in five proteins analyzed by PLIP [24] (S4 Table). For example, the predicted binding pose of the best AutoDock score conformation of PRKCB cognate docking was 2.79 Å when the MSM with TT structure was used as the receptor structure. However, out of eleven interacting residues identified in the reference structure, ten residues were retrieved in the MSM-ligand complex model. By analyzing the interaction pattern, the predicted docking pose to the multi-state modeled structure has hydrophobic interaction with L348, F353, V356, A369, K371, A483, and D484 and hydrogen bonding with T404, E421, and V423. These interactions are also observed in the crystal structure (S2 Fig). The cognate docking benchmark result would imply that MSM models could be used to predict binding poses of kinase-ligand complexes, thus they are suitable for virtual ensemble screening.

Similar trends are observed for the lowest RMSD conformations and average RMSD of 50 conformations. Among the AF2 predicted structures, MSM with TT shows the most accurate models (average of the lowest RMSD pose: 1.40 Å, success case: 19) and the standard AF2 performs slightly lower (average of the lowest RMSD pose: 1.50 Å, success case: 20). The docking accuracy of MSM with NT sets is the worst among the receptor structure sets (average of the lowest RMSD pose: 2.14 Å, success case: 14). In most cases, the best AutoDock score conformations do not match to the lowest RMSD conformations, which means that the AutoDock score could not be able to find the optimal docking poses. The average RMSD of 50 conformations follows the same order as the other two metrics: MSM with TT is the smallest, and MSM with NT is the highest.

### Virtual Screening Performance with Multi-state Models

To investigate the advantage of using the MSM technique for SBVS, the compound library for each kinase protein from DUD-E docked to structures with diverse states generated by using our method and compared to the structures modeled with standard AF2. AutoDock-GPU was used for the benchmark. For ensemble docking using generated models by MSM, since a molecule was docked to a couple of receptor structures and each compound-structure pair had a docking score, we needed to decide representative scores of the compounds to rank the molecules. We employed two types of representative scores: the AutoDock best (ADB) and the Boltzmann-weighted (BW) scores. The ADB score picks the lowest AutoDock score among the docked results, while the BW score is a weighted average of the scores across all structures. The details of the BW score are given in the Materials and Methods section.

Table 2 shows the performance of SBVS using the various receptor structure sets. To evaluate the performance, five metrics are used: enrichment factors at 1%, 5%, and 10% (EF_X%_), an area under ROC curve (AUC), and Boltzmann-enhanced discrimination of receiver operating characteristic (BEDROC). EF at X% indicates a capability of finding active molecules within the top X%, and AUC represents the discrimination power of a screening method between active and decoy molecules. BEDROC puts exponential weights to the early rank of molecules, thus it is able to solve the ‘early recognition problem’ caused in AUC [25]. Details of the metrics are illustrated in the Materials and Methods section. As observed in the cognate docking benchmark, MSM structures are better or equal to standard AF2 results, regardless of the scoring method or the template set for modeling. Details of individual results are provided in S5 Table. Comparing EF_1%_ values target-by-target, the MSM performed better than or equal to standard AF2 in 18 proteins out of 25 targets (72%) using the ADB score screened to the structures modeled with the NT set. Even with the lowest EF_1%_ combination, TT models with BW scoring, the MSM performed better than or equal to standard AF2 in more than half of proteins (13 proteins). Interestingly, although the TT set models with the same KinCoRe classification as the reference have more accurate structure and docking poses than the NT set models, the virtual screening performance is slightly worse in both scoring schemes.

**Table 2.** Performance of structure-based virtual screening on various receptor models.

|  | MSM with TT |  | MSM with NT |  | Standard |
| --- | --- | --- | --- | --- | --- |
|  | ADB | BW | ADB | BW | AF2 |

| DUD-E Kinase Subset (25 proteins) |  |  |  |  |  |
| --- | --- | --- | --- | --- | --- |
| **EF<sub>1%</sub>** | 7.2 | 7.2 | 8.2 | 8.0 | 6.6 |
| **EF<sub>5%</sub>** | 3.5 | 3.7 | 3.7 | 3.7 | 3.5 |
| **EF<sub>10%</sub>** | 2.7 | 2.7 | 2.6 | 2.7 | 2.6 |
| **AUC** | 0.660 | 0.639 | 0.661 | 0.664 | 0.659 |
| **BEDROC** | 0.193 | 0.196 | 0.183 | 0.189 | 0.183 |
| Average Dissimilarity $\geq 0.7$ (14 proteins) | | | | | |
| **EF<sub>1%</sub>** | 5.3 | 5.6 | 6.4 | 6.9 | 4.6 |
| **EF<sub>5%</sub>** | 3.0 | 3.1 | 3.2 | 3.2 | 2.8 |
| **EF<sub>10%</sub>** | 2.4 | 2.4 | 2.2 | 2.3 | 2.1 |
| **AUC** | 0.646 | 0.652 | 0.648 | 0.650 | 0.641 |
| **BEDROC** | 0.166 | 0.171 | 0.154 | 0.161 | 0.146 |
| Average Dissimilarity $< 0.7$ (11 proteins) | | | | | |
| **EF<sub>1%</sub>** | 9.6 | 9.3 | 10.6 | 9.3 | 9.1 |
| **EF<sub>5%</sub>** | 4.2 | 4.4 | 4.4 | 4.3 | 4.3 |
| **EF<sub>10%</sub>** | 3.0 | 3.1 | 3.1 | 3.1 | 3.3 |
| **AUC** | 0.678 | 0.681 | 0.678 | 0.681 | 0.681 |
| **BEDROC** | 0.228 | 0.228 | 0.220 | 0.226 | 0.229 |

To elucidate the variance in performance between MSMs using TT and NT models, we identified kinases that exhibited notable differences in EF_1%_ between the two sets. Of the kinases studied, both ABL1 and KDR demonstrated superior performance using the MSM with the NT set compared to the TT set, across both ensemble scoring methods (S5 Table). We measured TM-Scores of the individual models with the same structural state, those of templates used to predict the structures, and EF_1%_ of the models (S6 and S7 Tables). Among the kinase states analyzed, the DFGout-BBAminus for both proteins produced a significant difference in both templates and models. When the kinase forms an active state, the activation loop forms an extended conformation to facilitate the catalytic function of the protein, thus the DFGin conformation has a conserved structure. On the other hand, for the DFGout conformation, the activation loop collapsed on the protein surface with high flexibility [11]. Therefore, although the protein structures have the same KinCoRe notation with DFGout, the activation loop can have different structures. The TM-Scores of the templates to model the DFGout-BBAminus state are 0.88 and 0.96 for ABL1 and KDR, respectively. The structural difference of templates influenced the predicted models, resulting in TM-Scores 0.89 for both proteins. S3 Fig shows a structural difference of ABL1 predicted models with DFGout-BBAminus conformation. The difference of activation loop location also might affect to the virtual screening performance, leading TT set (3.28 and 8.78 for ABL1 and KDR, respectively) has lower EF_1%_ value than NT set (15.83 and 15.85 for ABL1 and KDR, respectively). Significantly, this difference of performance in the DFGout-BBAminus state impacted the overall ensemble docking results.

One benefit of using the MSM models is that the predicted models are diverse, so it is potentially useful for discovering various scaffolds of hit chemicals. To examine whether the hypothesis is true or not, we divided the DUD-E kinase subset into two based on the diversity of active compounds in the screening library. The pairwise Tanimoto coefficients (T_c_) between the active compounds were calculated by using RDKitFP fingerprint in RDKit [26]. Then the pairwise distances between the compounds are calculated as (1 – T_c_). The diversity of active compounds is defined as the average distances of the active compounds (S5 Table). With a threshold of 0.7, the kinases were classified into two groups, 14 proteins with higher than or equal to the cutoff and the remaining 11 targets.

For the kinases with more diverse active compounds, the ensemble screening with MSM models showed higher performance than standard AF2 models in all metrics. For EF_1%_, the MSM performed much better than standard AF2 (Table 2). Out of 14 proteins, the MSM with NT set and BW scoring has higher or equal EF_1%_ values than standard AF2 models in 11 targets. On the other hand, when the active compounds become less diverse, MSM still performed better than the standard AF2, but the gap between them declined for EF_1%_. This implies that our approach would be powerful for discovering diverse molecular scaffolds.

One of the diverse active hit examples is ABL1. The average dissimilarity of the active compounds is 0.73. Regardless of the template set and the scoring scheme, EF_1%_ of MSM ensemble screening showed higher performance (TT models with ADB: 9.3, TT models with BW: 10.4, NT models with ADB: 15.3, and NT models with BW: 17.5) than standard AF2 model (6.0). We also examined the diversity of active compounds ranked within the top 1% ranked molecules. Our ensemble protocol tends to find diverse hits: 0.51, 0.53, 0.63, and 0.62 for TT models with ADB, TT models with BW, NT models with ADB, and NT models with BW, respectively. In contrast, the diversity of active compounds using the standard AF2 is 0.27.

CSF1R is another example showing MSM ensemble screening is able to find diverse hit compounds. The average dissimilarity of hit molecules is 0.73. The gap of EF_1%_ between MSM models (5.4) and the standard AF2 model (3.6) is smaller than the case of ABL1. The diversities of discovered hit compounds within the top 1% are 0.70, 0.65, 0.73, and 0.73 in the order of TT models with ADB, TT models with BW, NT models with ADB, and NT models with BW, similar to the average dissimilarity of all active compounds. However, the hit compounds within the top 1 % identified using the standard AF2 model is 0.23. Fig 5 shows the docking pose of one of the active compounds, ChEMBL245377. The MSM ranked the active compound within the top 1% (10^th^ using TT models and BW scoring, out of 12316 compounds) which is not ranked by the standard AF2 structure (167^th^). The compound is not successfully docked using the the standard AF2 model.

**Fig 5.**
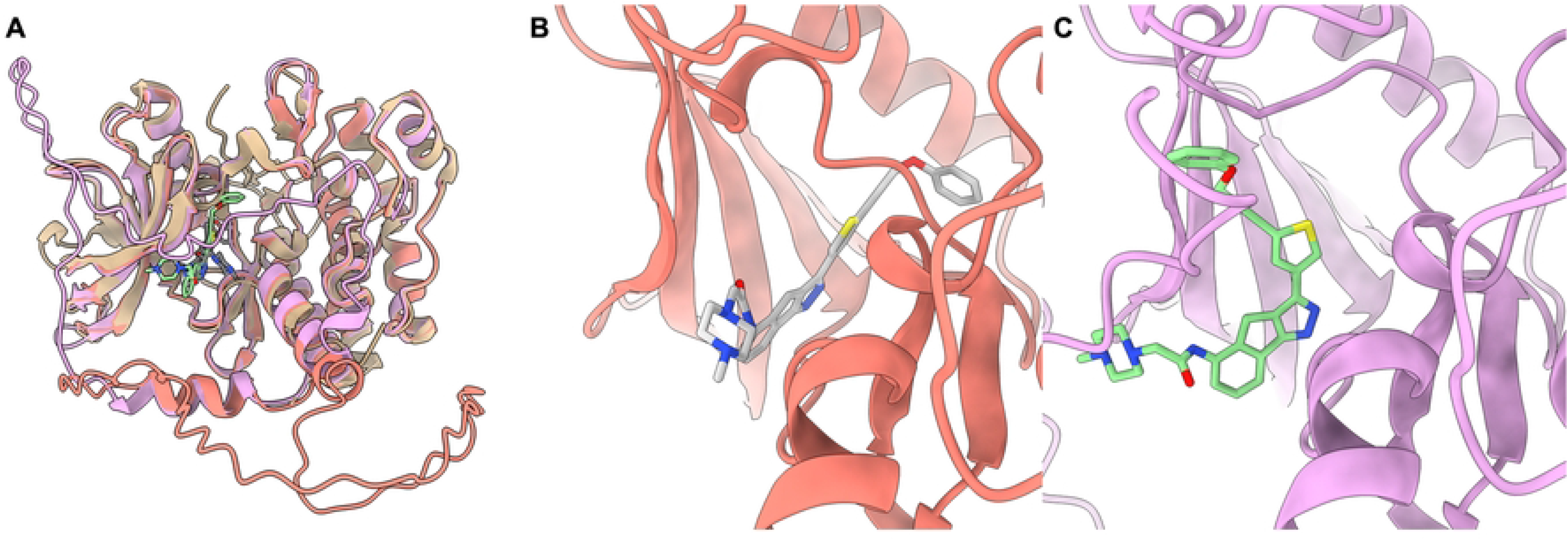
Predicted binding poses of ChEMBL245377 and CSF1R structures. **A**. The superimposed complex structures of ChEMBL245377 and CSF1R. The crystal structure of CSF1R (PDB ID:3KRJ), standard AF2 predicted structure, and MSM model (DFGout-BBAminus, with TT sets) are represented as ribbon diagrams colored as gold, pink, and orange, respectively, while the predicted docking poses of the compound are represented as sticks. **B,C**. Focused binding sites and docking poses of ChEMBL245377. The compound is ranked 10^th^ by ensemble docking (**B**) and 167^th^ in docking for the standard AF2 predicted structure (**C**), Although the ensemble screening with MSM structures performed generally better than standard AF2, there is an issue with selecting the representative score of a compound. For instance, for FGFR1, the MSM model with TT set and BW score showed EF_1%_ as 2.1. When we observed EF_1%_ values of individual models, the DFGin-BLBplus state outperformed than any other structures including standard AF2 (Table 3). Thus, a proper method for selecting or calculating representative scores for a compound should be designed to achieve high performance for MSM ensemble screening.

**Table 3.** EF_1%_ for FGFR1 structures of specific states. The MSM structures are built from TT set.

| Structural State | EF<sub>1%</sub> |
| --- | --- |
| DFGin-BLAminus | 2.9 |
| DFGin-BLAplus | 6.4 |
| DFGin-BLBminus | 0.7 |
| DFGin-BLBplus | 8.6 |
| DFGin-BLBtrans | 2.1 |
| DFGout-BBAminus | 3.6 |
| DFGout-Unassigned | 1.4 |
| MSM with BW score | 2.1 |
| Standard AF2 | 1.4 |

## Conclusion

The receptor conformation affects SBVS performance. Like other proteins, kinase adjusts the conformation of its binding site in response to the binding ligand. Therefore, it is crucial to have adequate kinase structure to obtain inhibitors with the required mode of action or diversity. For human kinase structures that were identified through experiments, however, there is a clear bias toward the active state. The prediction of the AF2 structure could be influenced by the bias in the PDB database. We noticed that there is a bias toward the active state in the predicted structures with standard AF2 protocol. The results of the SBVS using the predicted structure would be compromised by this bias in receptor structure. To overcome the bias, we applied the MSM technique by providing a structural template to AF2 to generate structures with diverse states and using the models for ensemble docking. Compared with standard AF2 models, MSM protocol produced more accurate or comparable models although it did not give MSA as an input. Also, in cognate docking study, MSM models provided ligand docking poses close to the crystal structure. With the diverse predicted kinase structure, we performed ensemble screening. The ensemble method showed enhanced or comparable SBVS performance to the standard AF2 modeled structure result. We also observed that our method would be more suitable when the ligands are diverse, leading to the identification of a diverse range of kinase inhibitors. Even for the targets that ensemble docking method could not find active compounds, we found that some of the models outperformed the standard AF2. Thus, the selection of a representative structure should be improved and remains the next work for this project.

Ensemble screening with MSM models would open the possibility of uncovering novel kinase inhibitors with diverse chemical scaffolds. It has advantages in addressing current challenges in kinase inhibitor development for finding chemically diverse compounds. The chemical diversity of kinase inhibitors could aid in overcoming the problem of drug resistance generally caused by the mutation, a significant obstacle in kinase-targeted cancer therapies. Additionally, it could increase chance to find hit compounds, not similar to the existing patents. By exploring a diverse array of kinase inhibitors with accurately predicted structures, we would be able to find inhibitors with different modes of action. This could lead to the development of novel therapeutic strategies that are more robust in the face of drug resistance. Hence, our approach could potentially be applied to more effective and precise kinase-targeted therapies. In addition, the MSM method could be applied to important therapeutic targets such as GPCRs and nuclear receptors.

## Materials and Methods

### Benchmark Dataset

To investigate whether our approach can generate a structural ensemble properly and improve SBVS performance or not, we selected a kinase subset of DUD-E [16]. For each target, a reference PDB structure for screening and a compound library composed of active and decoy molecules are provided in the DUD-E set. The active compounds of DUD-E set are composed of the molecules with affinity 1 μM or better were extracted from ChEMBL09 [5]. The reference structures were selected by considering the resolution and enrichment of finding active molecules by DOCK3.5. The decoys of the DUD-E set are constructed by gathering compounds with similar characteristic to the active molecules such as logP and number of rotatable bonds from ZINC database [27]. The kinase subset, which is used in this work, consists of 26 kinases with include 205.6 actives and 12,830 decoys on average. Among the 26 targets in DUD-E kinase subset, SRC kinase was removed from DUD-E benchmark set because the given reference structure is not originated from human.

### Kinase Structural State Annotation

In the active site of protein kinases, the activation loop, 20-30 residues long, is the most important secondary structural element [28] for determining the structural state. The loop starts from the conserved three-residue-long sequence, the DFG motif. In this work, the standalone version of KinCoRe [11] (https://github.com/vivekmodi/Kincore-standalone, Accessed 4/14/2022) was employed to annotate the conformational state of all experimental and modeled kinase structures. The program categorizes the conformational state into 12 classes by the location of the activation loop and dihedral angles of the DFG motif. The spatial state of the activation loop is defined by two distances: 1) a distance between Phe-ring of the DFG motif and Cα atom of the fourth residue from the conserved Glu in the C-helix of N-lobe and 2) a distance from the Phe-ring to the conserved Cζ atom of conserved Lys in β3 strand of N-lobe (Fig 1).

Based on the distances, the activation loop location is classified into three classes: DFGin, which is the Phe-ring located under C-helix, DFGout, the Phe-ring is moved into ATP binding pocket, and DFGinter, an intermediate state between DFGin and DFGout. The program further classifies the activation loop structural state by calculating dihedral angles: ϕ, ψ backbone dihedral angles of X-DFG (a residue before the DFG motif), Asp, and Phe of DFG motif, and χ1 angle of DFG-Phe. As a result, DFGin, the dominant class, has seven subclasses (BLAminus, BLAplus, ABAminus, BLBminus, BLBplus, BLBtrans, and Unassigned), while DFGinter (BABtrans and Unassigned) and DFGout (BBAminus and Unassigned) have only two subclasses. The three letters after the activation loop states follows the region of Ramachandran map occupied by X, D, and F residues: A, B, L for alpha, beta, and left-handed, respectively. The χ1 angle of Phe is indicated as plus (+60 degree), minus (−60 degree), and trans (180 degree). The last class is Unassigned-Unassigned, the activation loop and DFG conformations cannot be determined.

### Construction of Template Database for Each Structural State

To construct the structural template database for MSM, KLIFS [18], a database of experimentally determined kinase structures was used. The database contains catalytic domain structures of kinases, extracted from PDB and their inhibitors and provides the interaction information between the protein and the compound. As of May 2023, the database is composed of 6,344 structures (13,382 monomers). Among the kinase structures in KLIFS (Accessed 1/18/2023), we filtered out the non-human proteins and proteins that produced errors during KinCoRe annotation, resulting in 11,106 monomer structures. To construct the template structure database for each state, the crystal structures with the same annotation by KinCoRe were gathered.

### Standard AlphaFold2 Modeling

The kinase structure was modeled with the standard protocol of AF2 (v.2.3.1) to compare with MSM models. Only the kinase domain was modeled from the full sequence of a protein. The MSA for the kinase sequence was generated from BFD (v.3.2019), MGnify (v.5.2022), and UniRef90 (v.2.2022) via HHblits (from HH-suite v3.3.0) and Jackhammer (from HMMER v3.3.2). Four highest sequence identity proteins with 3D atomic coordinates were selected to provide template structures. The number of recycles was set to three and the model relaxation step was integrated into the procedure. As AF2 has five different trained models and runs all of them independently in a single run, five structures were generated from a single run. The models with the highest plDDT score out of the five predicted structures for the comparison, since plDDT is a confidence measure of AF2 predicted models.

### Multistate Modeling of Kinase using Structural Template

The workflow of MSM is given in Fig 3. From a given target kinase sequence to be modeled, the templates were searched by MMseqs2 (release 11) easy-search (e-value cutoff: 1e-3) [29] against all sequences in each structural state. For each structural state, the top five templates, which were determined by e-value, were used for the modeling. To mimic the real drug discovery process and check the influence of the template for virtual screening, we generated two template sets for modeling: one set has the query protein which means 100% sequence identity with the query sequence (TT set), and the other template sets without the query protein (NT set). Modeling with each template was conducted independently.

Since AF2 produced five models per single run, 25 structures for each specific state were generated in total. Among them, we selected one structure for benchmarking with two criteria: conformational state and quality of the predicted model. First, we filtered out the models with different KinCoRe annotations from the template structure classification. For example, to predict a model the DFGin-ABAminus conformation of ABL1, five templates structure of TYR family (LYN: 5XY1, EPHA2: 7KJB, 5NK3, 4TRL, and IGF1R: 3F5P) were selected. However, the models were annotated as DFGin-BLAminus rather than DFGin-ABAminus conformation (S4 Fig). Thus, all predicted models were discarded. After filtering by the KinCoRe annotation, the models with plDDT less than 70 were also removed. Among the remained structures, the model with the highest plDDT score was finally selected. Other details for modeling are the same as standard AF2 modeling.

### Assessment of Model Quality

The TM-score [19] evaluates the structural similarity of protein structures. It is scaled according to the size of the protein and exhibits a better sensitivity to the overall structural alignment compared to the RMSD. The models with the same KinCoRe notation as the crystal structures, extracted from the DUD-E set were chosen for comparison with the crystal structure employing the TM-score. The TM-score’s values range from 0 to 1, where 1 signifies a perfect alignment.

MolProbity [17] is a comprehensive validation tool for the structural integrity of proteins and nucleic acids. MolProbity validates protein structures through hydrogen replacement, comprehensive all-atom contact analysis, and evaluation of torsional angles. The MolProbity score integrates various factors into a single metric to indicate the model’s reliability, where a lower score denotes a better model. The models from the standard AF2 and MSM were validated using MolProbity implemented in Phenix software [30].

### Docking and Virtual Screening using AutoDock-GPU

AutoDock-GPU 1.5.3 [20], which is open-source and GPU-accelerated, was employed to benchmark the virtual screening performance of kinase structures. We used AutoDockTools [31] to convert the receptor PDB files to PDBQT and Meeko [32] based on RDKit [26] to convert the ligand files into PDBQT format.

To define a docking pocket location, the model structure was superimposed with the reference crystal structures provided in the DUD-E set using PyMOL alignment module [33]. Then, a cubic box centered at the geometrical center position of the cognate ligand structure of the reference protein structure was defined. Each dimension of the box has a size of 22.5 Å, a default option of the program. The parameter nrun, the number of pose generations and searches in AutoDock-GPU, was set to 50 to find the optimal AutoDock score between protein and ligand.

### Scoring Schemes for Ensemble Docking

To get the representative score of a ligand that docked to the multiple receptor structures in ensemble screening, we employed two scoring schemes: AutoDock best score (ADB) and Boltzmann-weighted score (BW). After gathering all AutoDock scores of a compound docked to the MSM structures, ADB scheme picks the lowest value as a representative score for the ligand. For example, if a ligand is docked to five kinase structures with docking scores of −11 kcal/mol, −12 kcal/mol, −8 kcal/mol, −7 kcal/mol, and −9 kcal/mol, then the compound has a score of −12 kcal/mol.

Instead of using the docking score from a single structure, the BW scheme calculates a weighted average of the docking scores. We modified BW score of Shin et al. [1], which was originally used to calculate a score of a protein with multiple ligand conformations, to apply a single ligand to multiple protein conformations (Equation 1).

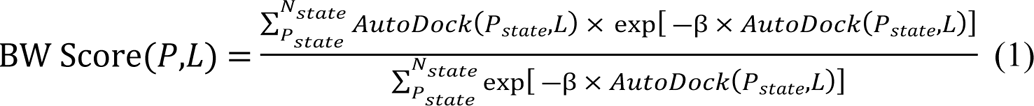

where *β* = 1, P and L are protein and ligand, respectively. P_state_ means target protein structure with a specific state. AutoDock(P_state_, L) means AutoDock score of a ligand for the target protein with a specific state.

### Evaluation Metrics for Docking and Screening

To evaluate the performance of cognate docking, RMSDs of the docked conformations from the bound conformation of the crystal structure were calculated. Then we picked two conformations: one with lowest AutoDock score and the other one is the lowest RMSD conformation. We also measured the docking success rate of the 24 target proteins with the RMSD cutoff of 2 Å, a widely used criterion for many docking studies [21–23]. In order to compare the virtual screening performance of MSM model ensemble screening with X-ray crystallography and standard AF2 structures, the EF_X%_, AUC, and BEDROC were calculated.

The EF_X%_ is a widely used metric to evaluate virtual screening methods. The enrichment factor quantifies the extent to which active compounds are sampled in the top N% of compounds relative to the total compound set (Equation 2).

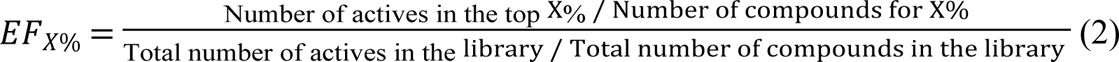

We set X as 1, 5, and 10. A random selection of compounds makes the EF value 1. One of the most popular measures for the discrimination problem is AUC. The true positive rate in relation to the false positive rate was plotted to create a receiver operating characteristic curve. In the case of virtual screening, the ratio of active chemicals represents the true positive rate, while the ratio of decoy molecules represents the false positive rate. When a program detects all active compounds before ranking any decoy compounds, AUC reaches 1.0, the maximum value and AUC 0.5 means that the program performance is the same as the random selection. Although AUC gives an overall performance discriminating power between actives and decoys of SBVS, it has a problem called ‘early recognition’ [25]. In virtual screening, the highly ranked compounds are passed to experiment, not all compounds. Thus, it is important to rank active molecules within high rank. To solve this issue, Boltzmann-enhanced discrimination of receiver operating characteristic (BEDROC) puts an exponential weight on the highly ranked active compounds (Equation 3).

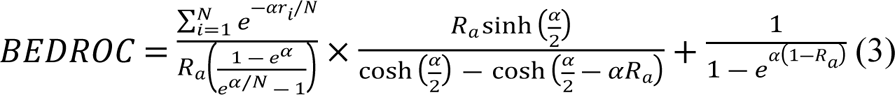

N is the number of compounds, R_a_ is the ratio of active compounds in the library, i the index of the active compounds, and r_i_ is the rank of the active compound i. In this work, the weight, α, is set to 20.

## Acknowledgements

All authors acknowledge the support from the Bio & Medical Technology Development Program of the National Research Foundation (NRF) funded by the Korean government (No. 2022M3E5F3081268). WHS also acknowledges support from Korea University Grant (No. K2327351). JL also acknowledges the support from NRF Grants funded by the Korean government (MSIT) (Nos. 2022R1C1C1005080 and 2020M3A9G7103933) and Korea Environment Industry & Technology Institute (KEITI) through “Advanced Technology Development Project for Predicting and Preventing Chemical Accidents” Program, funded by Korea Ministry of Environment (MOE) (RS-2023-00219144).

## Supporting Information

**S1 Table.** Percentage of human kinase structural states in PDB database, standard AlphaFold2, and average plDDT and MolProbity of predicted structures by multi-state modeling protocol.

**S2 Table.** The number of predicted models generated by MSM protocol.

**S3 Table.** Cognate docking result of 24 DUD-E kinase proteins. The values out of the parentheses are RMSD (Å) of the best AutoDock score while in the parentheses are the conformation of the lowest RMSD.

**S4 Table.** Percentage of retrieved interacting residues of the crystal structures from the failed cognate MSM docking benchmark cases.

**S5 Table.** Virtual screening results of individual proteins.

**S6 Table.** Details in multi-state modeling of ABL1.

**S7 Table.** Details in multi-state modeling of KDR.

**S1 Fig.** Distribution of MolProbity score by modeling method. The average MolProbity score was 1.04, 1.21, 1.24 for AF2, TT and NT respectively.

**S2 Fig.** Interacting residues identified by PLIP from the crystal structure of PRKCB (PDB ID: 2IOE, **A**) and docking results with the MSM model with TT (**B**). The interacting residues are colored as blue while the docked ligands are shown in yellow. The RMSD between the conformation is 2.79 Å. The hydrophobic interaction between the molecules is represented as dashed lines while the hydrogen bonding is shown as blue solid lines.

**S3 Fig.** Predicted structures of ABL1 with DFGout-BBAminus state. TT model (template: 7HZ0) is shown in sky blue and NT model (template: 3PYY) is shown in gold. The activation loop of each model is colored as green and orange for TT model and NT model, respectively.

**S4 Fig.** Example of modeling result that the conformational state is not matched with template. **A.** Overall structure comparison between the modeled structure (colored in beige) and the template (PDB ID: 5XY1, colored in sky blue). **B.** Focused view for the binding site. **C.** Ramachandran plot for modeled structure and the template. The modeled structure were annotated as DFGin-BLAminus (red x), while the template was DFGin-ABAminus (green x).

